# Genome-wide analysis yields new loci associating with aortic valve stenosis

**DOI:** 10.1101/213595

**Authors:** Anna Helgadottir, Gudmar Thorleifsson, Solveig Gretarsdottir, Olafur A. Stefansson, Vinicius Tragante, Rosa B. Thorolfsdottir, Ingileif Jonsdottir, Thorsteinn Bjornsson, Valgerdur Steinthorsdottir, Niek Verweij, Jonas B. Nielsen, Wei Zhou, Lasse Folkersen, Andreas Martinsson, Mahyar Heydarpour, Siddharth Prakash, Gylfi Oskarsson, Tomas Gudbjartsson, Arnar Geirsson, Isleifur Olafsson, Emil L. Sigurdsson, Peter Almgren, Olle Melander, Anders Franco-Cereceda, Anders Hamsten, Lars Fritsche, Maoxuan Lin, Bo Yang, Whitney Hornsby, Dongchuan Guo, Chad M. Brummett, Gonçalo Abecasis, Michael Mathis, Dianna Milewicz, Simon C. Body, Per Eriksson, Cristen J. Willer, Kristian Hveem, Christopher Newton-Cheh, J. Gustav Smith, Ragnar Danielsen, Gudmundur Thorgeirsson, Unnur Thorsteinsdottir, Daniel F. Gudbjartsson, Hilma Holm, Kari Stefansson

## Abstract

Aortic valve stenosis (AS) is the most common valvular heart disease, characterized by a thickened and calcified valve causing left ventricular outflow obstruction. Severe AS is a significant cause of morbidity and mortality, affecting approximately 5% of those over 70 years of age^1,2,3^. Little is known about the genetics of AS, although recently a variant at the *LPA* locus^4^ and a rare *MYH6* missense variant were found to associate with AS^5^. We report a large genome-wide association study (GWAS) with a follow-up in up to 7,307 AS cases and 801,073 controls. We identified two new AS loci, on chromosome 1p21 near *PALMD* (rs7543130; OR=1.20, *P*=1.2×10^−22^) and on chromosome 2q22 in *TEX41* (rs1830321; OR=1.15, *P*=1.8×10^−13^). Rs7543130 also associates with bicuspid aortic valve (BAV) (OR=1.28, *P*=6.6×10^−10^) and aortic root diameter (*P*=1.30×10^−8^) and rs1830321 associates with BAV (OR=1.12, *P*=5.3×10^−3^ and coronary artery disease (CAD) (OR=1.05, P=9.3×10^−5)^. These results indicate that AS is partly rooted in the same processes as cardiac development and atherosclerosis.

---

We tested 32.5 million sequence variants for association with AS in 2,457 Icelandic cases and 349,342 controls. The variants were identified by whole-genome sequencing 15,220 Icelanders and imputing into 151,678 chip-typed, long-range phased individuals and their close relatives^6^. We observed one genome-wide significant association between AS and the intergenic variant rs7543130 (risk allele frequency (RAF) [A]=51.2%) on chromosome 1p21 near the *PALMD* gene (OR=1.23; 95% confidence interval (CI): 1.15-1.31, *P*=6.8×10^−10^ (significance threshold for intergenic variants set at *P*=7.9×10^−10^, see Methods and ref.^7^)). We noted that rs7543130 was recently reported to associate with aortic root size in an independent GWAS^8^ and we replicate this association in our Icelandic aortic root dimension sample (*P*=1.30×10^−8^) (Table 2).

We tested rs7543130 and six additional variants that did not reach the significance threshold in up to 4,850 AS cases and 451,731 controls from Sweden, Norway, United Kingdom and the United States (Table 1, Supplementary Table 1). The joint analysis showed robust association between AS and rs7543130 (OR=1.20; 95% CI: 1.16-1.25; *P*=1.2×10^−22^). In addition, the rs1830321 variant (RAF[T]=37.5%) intronic to *TEX41,* a non-protein coding gene on chromosome 2q22, associated genome-wide significantly with AS in the joint analysis (OR=1.15; 95% CI: 1.11-1.20, *P*=1.8×10^−13^) (Table 1).

**Table 1.** Meta-analysis results for aortic stenosis variants

|  | Cases/controls | *PALMD* intergenic<br>rs7543130 [A/C]<br>EAF=51.2% |  | *TEX41* intronic<br>rs1830321 [T/C]<br>EAF=37.5% |  | *LPA* intronic<br>rs10455872 [G/A]<br>EAF=6.2% |  |
| --- | --- | --- | --- | --- | --- | --- | --- |
|  |  | OR (95% CI) | *P* | OR (95% CI) | *P* | OR (95% CI) | *P* |
| Iceland | 2,457/349,342 | 1.23 (1.15-1.31) | $6.8 \times 10^{-10}$ | 1.20 (1.12-1.28) | $7.6 \times 10^{-8}$ | 1.4 (1.23-1.56) | $1.8 \times 10^{-7}$ |
| Sweden (MDCS)* | 470/15,162 | 1.14 (0.98-1.33) | 0.092 | 1.21 (1.05-1.39) | 0.0080 | 1.55 (1.28-1.88) | $1.0 \times 10^{-5}$ |
| Sweden, Stockholm | 318/1,376 | 1.25 (0.98-1.59) | 0.068 | 1.18 (0.89-1.56) | 0.24 | 1.19 (0.90-1.57) | 0.23 |
| UK-Biobank | 1,844/406,814 | 1.25 (1.17-1.33) | $3.8 \times 10^{-11}$ | 1.14 (1.06-1.22) | $1.36 \times 10^{-4}$ | 1.54 (1.38-1.71) | $4.8 \times 10^{-15}$ |
| Norway (HUNT) | 1,546/24,235 | 1.13 (1.05-1.22) | 0.0012 | 1.11 (1.02-1.20) | 0.010 | 1.48 (1.28-1.71) | $1.0 \times 10^{-7}$ |
| USA, Michigan | 251/2,510 | 1.15 (0.96-1.39) | 0.13 | 1.01 (0.85,1.24) | 0.76 | 1.32 (0.94-1.84) | 0.10 |
| **Combined** | **6,886/799,439** | **1.20 (1.16-1.25)** | **$1.2 \times 10^{-22}$** | **1.15 (1.11-1.20)** | **$1.8 \times 10^{-13}$** | **1.46 (1.37-1.56)** | **$1.9 \times 10^{-31}$** |

Results are shown for the discovery and follow-up data sets and the joint analysis (combined). The effect allele is the first allele in brackets [effect allele/non-effect allele]. The EAF (effect allele frequency) is for the Icelandic population. OR, allelic odds ratio; 95% CI, 95% confidence interval; *P*, *P* value from case control association analysis. \* The association results for the rs10455872 variant in the MDCS (Malmö Diet and Cancer study) included 613 cases and 28,109 controls.

We replicate the reported association of the intronic *LPA* variant^4^ rs10455872 with AS in Iceland and the follow-up sample sets (combined OR=1.46; 95% CI: 1.37-1.56, *P*=1.9×10^−31^) (Table 1). AS is usually graded as mild, moderate, or severe, based on the valve area and the pressure gradient across the valve. The estimated 5-year survival in symptomatic severe AS ranges from 15% to 50%^3^ unless outflow obstruction is relieved by aortic valve replacement, the only definitive treatment for AS. We specifically tested the association of the two novel AS variants and the *LPA* variant, with a subset of Icelandic AS cases who had undergone aortic valve replacement, representing those with severe AS. The effect sizes were not significantly different from those for all AS (Supplementary Table 2).

Although the underlying pathophysiology of AS is not fully understood, studies show that calcified aortic valve lesions have many characteristic features of atherosclerosis, including initial endothelial damage, oxidized lipid deposition, chronic inflammation and calcification^9^. In addition it is recognized that BAV, the most common congenital cardiac malformation, accelerates the development of AS by decades^9^. BAV is found in up to half of those with severe AS^10^ while the prevalence of BAV is 0.5-2% in the general population. A rare p.Arg721Trp *MYH6* missense variant (RAF=0.34%) that was previously shown to associate with cardiac arrhythmia^11,12^, has recently been reported to also associate with coarctation of the aorta, risk of BAV (OR=8.04; 95% CI: 3.36-19.22; *P*=2.8×10^−6^) and with AS (OR=2.65; 95% CI: 1.78-3.96; *P*=1.8×10^−6^) (ref.^5^). For the *MYH6* variant, the effect on BAV risk is substantially greater than that for AS (P=0.023), suggesting that the AS risk conferred by this variant is mediated through BAV. We tested rs7543130 near *PALMD,* rs1830321 in *TEX41,* and the *LPA* rs10455872 for association with BAV in 1,555 cases and 33,883 controls from Iceland, Sweden and the United States. Both novel AS variants associated with BAV (OR=1.28; 95% CI: 1.19-1.39; *P*=6.6×10^−10^ for rs7543130 and OR=1.12, 95% CI: 1.04-1.22, *P*=5.3×10^−3^ for rs1830321). The LPA rs10455872 did not associate with BAV (Table 2).

**Table 2.** Association of aortic stenosis variants with other cardiovascular traits

| Phenotype (cc) | Cases/controls | PALMD intergenic<br>rs7543130 [A/C] |  | TEX41 intronic<br>rs1830321 [T/C] |  | LPA intronic<br>rs10455872 [G/A] |  | MYH6 missense<br>p.Arg721Trp [A/G] |  |
| --- | --- | --- | --- | --- | --- | --- | --- | --- | --- |
|  |  | OR (95% CI) | P | OR (95% CI) | P | OR (95% CI) | P | OR (95% CI) | P |
| **Bicuspid aortic valve (BAV)** | 208/25,139 | 1.26 (0.99, 1.60) | 0.059 | 1.31 (1.03, 1.67) | 0.025 | 1.12 (0.72, 1.75) | 0.61 | 8.04 (3.36, 19.22) | 2.8×10<sup>-6</sup> |
| Sweden-Stockholm | 275/1,516 | 1.27 (1.06, 1.52) | 0.0098 | 1.20 (0.99, 1.46) | 0.063 | 1.02 (0.92, 1.12) | 0.77 |  |  |
| USA-Houston | 147/864 | 1.29 (0.99, 1.67) | 0.057 | 1.25 (0.97, 1.60) | 0.085 | 0.96 (0.51, 1.81) | 0.91 |  |  |
| USA-Boston | 452/1,634 | 1.27 (1.09, 1.48) | 0.002 | 1.10 (0.95, 1.28) | 0.21 | 1.44 (1.08, 1.93) | 0.014 |  |  |
| USA-Michigan | 473/4,730 | 1.31 (1.14, 1.51) | 1.2×10<sup>-4</sup> | 1.00 (0.86, 1.16) | 0.97 | 1.19 (0.93, 1.54) | 0.17 |  |  |
| Combined BAV | 1,555/33,883 | 1.28 (1.19, 1.39) | 6.6×10<sup>-10</sup> | 1.12 (1.04, 1.22) | 5.3×10<sup>-3</sup> | 1.07 (0.98, 1.16) | 0.13 |  |  |
| **Atrial septal defect** | 708/353,019 | 1.23 (1.07, 1.42) | 3.9×10<sup>-3</sup> | 1.22 (1.06, 1.41) | 5.9×10<sup>-3</sup> | 1.15 (0.87, 1.52) | 0.32 | 3.17 (1.47, 6.81) | 3.2×10<sup>-3</sup> |
| **Ventricular septal defect** | 902/357,428 | 1.23 (1.07, 1.42) | 4.8×10<sup>-3</sup> | 1.04 (0.90, 1.21) | 0.59 | 1.14 (0.84, 1.53) | 0.41 | 4.40 (2.14, 9.07) | 5.7×10<sup>-5</sup> |
| **Coronary artery disease** | 37,782/318,845 | 1.00 (0.97, 1.02) | 0.74 | 1.05 (1.03, 1.08) | 9.3×10<sup>-5</sup> | 1.28 (1.21, 1.34) | 2.4×10<sup>-22</sup> | 1.21 (1.00, 1.48) | 0.056 |
| **Phenotype (qtl)** | **N** | **β (SE)** | **P** | **β (SE)** | **P** | **β (SE)** | **P** | **β (SE)** | **P** |
| **Aortic root diameter** | 19,513 | 0.065 (0.01) | 1.3×10<sup>-8</sup> | -0.017 (0.02) | 0.16 | -0.052 (0.02) | 0.020 | -0.068 (0.08) | 0.40 |

Association of aortic stenosis variants with several cardiovascular phenotypes in Icelandic datasets. Follow-up and joint analysis (combined BAV) is also provided for the association with bicuspid aortic valve (BAV). The effect allele is the first allele in brackets [effect allele/non-effect allele]. Cc, case-control. Qtl, quantitative trait. OR, allelic odds ratio; 95% CI, 95% confidence interval; *P*, *P* value. The rs-name for *MYH6* missense p.Arg721Trp is rs387906656. The effect (β) for aortic root diameter is given in standardized units. SE, standard error.

Next we examined the association of the AS variants with other cardiovascular diseases in the deCODE genetics phenotype database (Table 2 and Supplementary Table 2). In line with the association of p.Arg721Trp in *MYH6,* rs7543130, and rs1830321 with congenital BAV, all three associated with ventricular-and/or atrial septal defects, suggesting that these variants affect cardiac developmental pathways and through them, at least in part, the pathogenesis of AS. Rs1830321 in *TEX41* associates with CAD in Iceland (OR=1.05, 95% CI: 1.03-1.08; *P*=9.3×10^−5^), but not the *MYH6* missense variant or rs7543130 near *PALMD* (Table 2, Supplementary Table 2). The *TEX41* rs1830321 is in linkage disequilibrium (LD) with a known GWAS CAD variant rs2252641 at the same locus (R^2^=0.80)^13^. The LPA variant rs10455872 like rs1830321 also associates with CAD in addition to AS. This indicates that these variants influence atherosclerosis-like processes involving the aortic valve.

Several atherosclerotic risk factors associate with risk of AS in epidemiological studies^15^ and genetic predisposition to both elevated lipoprotein (a) (Lp(a)) and LDL-cholesterol, established risk factors of CAD, have been associated with risk of AS^4,16^. We thus tested the novel AS variants for association with traditional cardiovascular risk factors and found nominally significant association between rs1830321 and systolic and diastolic blood pressure in Iceland (Supplementary Table 3) and in data from the UK-Biobank (https://biobankengine.stanford.edu/search#).

The frequent comorbidity between AS and CAD^17^ together with the similarities in pathophysiology^9^, is consistent with shared genetic risk factors. To examine sharing of genetic predisposition, beyond the previously reported LPA variant and the rs1830321 on chromosome 2q22 reported in this study, we tested seventy-one other CAD variants^18,19^ for association with AS, both individually (Supplementary Table 4) and as a weighted genetic risk score (GRS) (Table 4).

In the Icelandic and UK-Biobank datasets combined, four CAD variants associate with AS at a significance threshold set at *P* = 7.0×10^−4^ = 0.05 / 71. These are the LPA variant rs3798220 (p.Ile1891Met), rs116843064 in *ANGPTL4* (p.Glu40Lys), rs646776 at the *CELSR2 / PSRC1* locus, and rs3184504 in *SH2B3* (p.Trp60Arg) (Table 3, Supplementary Tables 4 and 5). Apart from rs3184504 in *SH2B3* that is known to affect various phenotypes^20^, these variants affect either Lp(a)^14^, or non-HDL-cholesterol levels^21,22^.

**Table 3.**
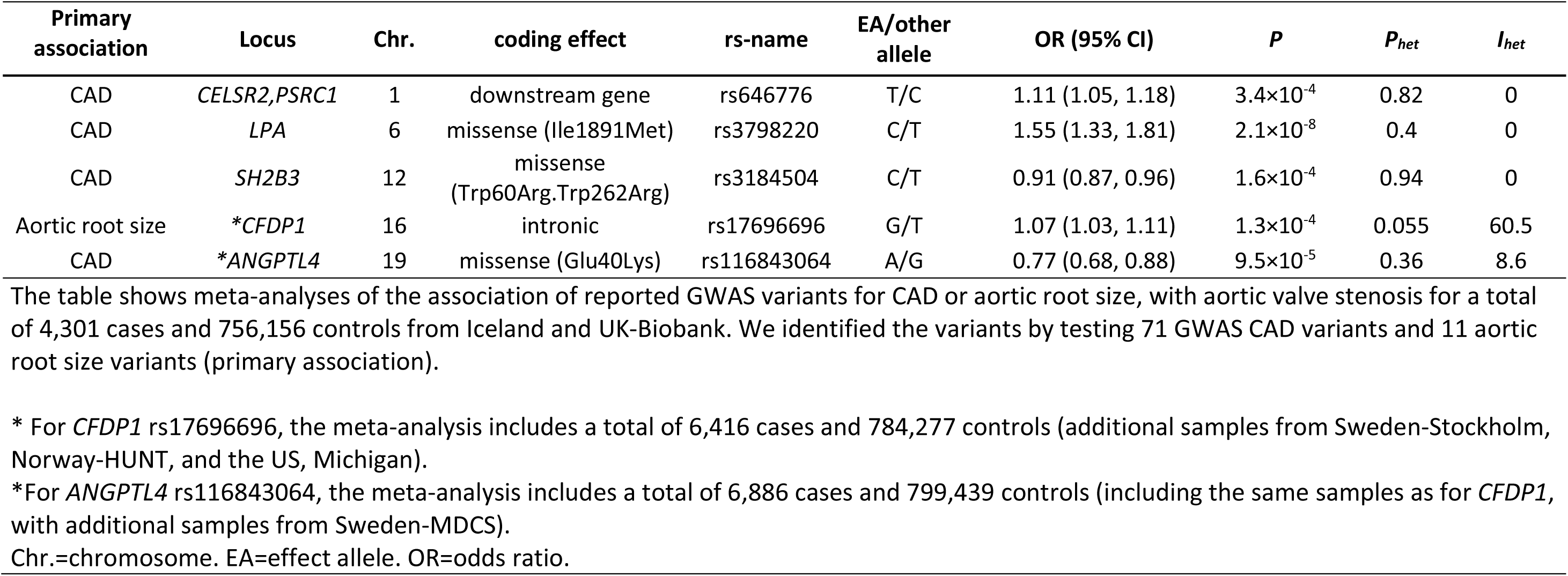
Aortic valve stenosis associations identified in analyses of reported CAD and aortic root size variants

Consistent with a shared genetic risk, the CAD-GRS associates with AS both in the Icelandic and the UK-Biobank datasets (combined *P*=7.5×10^−9^) (Table 4). However, the effect on AS was only around 37% of the effect on CAD and this association was no longer significant after adjustment for CAD diagnosis. Given the reported association between genetic predisposition to both elevated Lp(a) and LDL-cholesterol and AS^4,16^, we tested a subset of CAD-GRS (labelled as CAD-GRS-*lip*), constructed based on fourteen LPA and LDL-/non-HDL-cholesterol variants (Supplementary Table 4) for association with AS. CAD-GRS-*lip* associated strongly with AS *P*=5.1×10^−19^) with effect that was similar or larger than that for CAD (β = 0.91 and 1.02, for CAD and AS, respectively). This association with AS remained after adjusting for CAD diagnosis (*P*=5.1×10^−9^) (Table 4), suggesting that the genetic predisposition to elevated Lp(a) and LDL-cholesterol explains in large part the comorbidity between CAD and AS. Specifically, examining the impact of CAD-GRS-*lip* on the risk of AS among CAD cases shows that CAD cases with genetic predisposition to high Lp(a) or LDL-/non-HDL-cholesterol are at a greater risk of having AS, than CAD without such predisposition (*P*= 8.5×10^−7^) (Supplementary Table 6). In contrast, the complementary subset of CAD-GRS in which the *LPA* and LDL-/non-HDL-cholesterol variants are excluded, associated with less risk of AS, after adjusting for CAD status (*P* =0.0036) (Table 4). Given the known association of rs7543130 on chromosome 1p21 with aortic root dimension^8^ and the recognized relationship between BAV, AS, and aortopathy^23,10^, we tested eleven other reported aortic root size variants^8^ for association with AS. One of these variants, rs17696696 intronic to *CFDP1,* showed suggestive association with AS in the Icelandic and UK-Biobank datasets (Supplementary Table 7, Table 3). A correlated variant rs4888378 (R^2^=0.98) has been reported to associate with risk of CAD^24^, an association that is also observed in Iceland (OR=1.05, 95% CI: 1.03-1.07, *P*=4.5×10^−5^). None of the other AS variants associate with aortic root size (Table 2).

**Table 4.** The association of CAD genetic risk score with aortic valve stenosis

|  |  | Iceland |  | UK-Biobank |  | Iceland and UK-Biobank combined |  |
| --- | --- | --- | --- | --- | --- | --- | --- |
| | | $\beta$ | $P$ | $\beta$ | $P$ | $\beta$ (95% CI) | $P$ |
| **CAD-GRS-All** | **CAD** | 0.77 | $5.2 \times 10^{-153}$ | 0.76 | $< 10^{-300}$ | 0.76 (0.73,0.79) | $< 10^{-300}$ |
| . | **AS** | 0.29 | 0.00027 | 0.28 | $7.2 \times 10^{-6}$ | 0.28 (0.19,0.38) | $7.5 \times 10^{-9}$ |
| . | **AS<sub>adjCAD</sub>** | 0.03 | 0.62 | -0.05 | 0.45 | -0.01 (-0.10,0.08) | 0.83 |
| **CAD-GRS-lip** | **CAD** | 0.89 | $1.4 \times 10^{-36}$ | 0.92 | $7.8 \times 10^{-99}$ | 0.91 (0.84-0.99) | $1.4 \times 10^{-134}$ |
| . | **AS** | 1.05 | $2.3 \times 10^{-9}$ | 0.99 | $3.8 \times 10^{-11}$ | 1.02 (0.79-1.24) | $5.1 \times 10^{-19}$ |
| . | **AS<sub>adjCAD</sub>** | 0.77 | $1.5 \times 10^{-5}$ | 0.60 | $6.6 \times 10^{-5}$ | 0.67 (0.45,0.90) | $5.1 \times 10^{-9}$ |
| **CAD-GRS-non-lip** | **CAD** | 0.73 | $6.3 \times 10^{-120}$ | 0.72 | $4.3 \times 10^{-293}$ | 0.73 (0.69-0.78) | $< 10^{-300}$ |
| . | **AS** | 0.14 | 0.074 | 0.14 | 0.048 | 0.14 (0.04-0.24) | 0.0076 |
| . | **AS<sub>adjCAD</sub>** | -0.11 | 0.15 | -0.18 | 0.0088 | -0.15 (-0.25,-0.05) | 0.0036 |

GRS (Genetic Risk Score). CAD (coronary artery disease). AS (aortic valve stenosis).
CAD-GRS-All is based on 71 reported CAD variants. CAD-GRS-lip is based on 14 CAD variants with reported association with LDL-cholesterol (or non-HDL-cholesterol) or variants at the *LPA* locus. GRS-non-lip is based on 57 CAD variants, the same as in CAD-GRS-All, but excluding variants in CAD-GRS-lip). The effects of the GRSs on CAD are shown for comparison.
Number of cases/controls in Iceland and UK-Biobank respectively: CAD = 17,488/124,620 and 26,384/382,294; AS = 1,591/140,517 and 1,844/406,814.

Attempting to find causal variants and genes at the *PALMD* and *TEX41* loci, we looked for associations with expression quantitative trait loci (eQTL) using the Genotype-Tissue Expression (GTEx) dataset^25^, but did not find significant eQTL in 44 diverse human tissues from adults. Potential explanations for the lack of eQTL association could include: insufficient power to detect true associations with genes characterized by low expression, such as non-coding RNAs; the effect is specific to particular tissue type not represented in GTEx; or the impact of the variants might only emerge during development, but not in adult tissue.

To further investigate potential functional relevance of the two AS loci we mapped variants in LD (R^2^>0.5) with the lead variants to regulatory regions in heart and aorta tissue samples using public data from the NIH Roadmap Epigenomics Consortium^26,27^. Subsequently we used chromatin interaction maps^28^ for aorta, and left-and right-ventricle tissue samples to look for interactions between the regulatory regions, to which AS risk variants mapped, and promoters. At the *PALMD* locus, four variants mapped to regulatory regions annotated as enhancers and poised promoter (Figure 1-A, upper panel). Multiple chromatin interactions were observed for the regulatory regions harboring these four variants in left ventricle (Supplementary Figure). Notably, only the region harboring rs1890753 (R^2^ =0.97 with rs7543130) interacted with gene promoters (Figure 1-A, lower panel), including the promoters of *PALMD, SNX7, PLPPR5, PLPPR4,* and the non-coding RNAs, *LOC100129620* and *LOC101928270.* Thus, at the *PALMD* locus rs1890753 represents the candidate causal variant. In fetal heart tissue, a “poised” state is found at the rs1890753 locus (Fig 1-A upper panel). Poised chromatin is defined by the presence of histone modifications associated with both gene activation and repression, and is considered to be involved in the expression of developmental genes. This observation is in line with an impact of the rs1890753 risk variant during fetal development.

**Figure 1.**
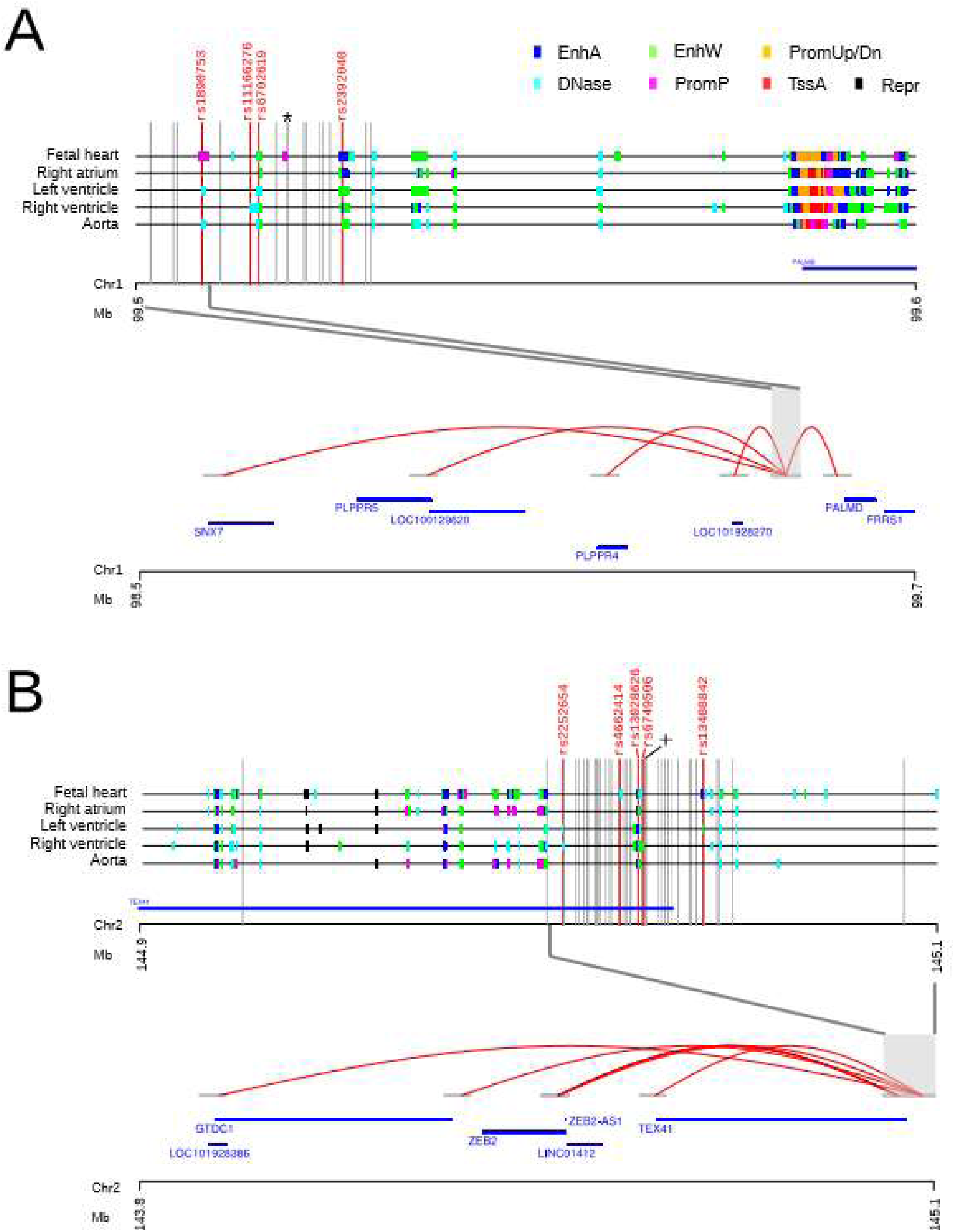
Chromatin states indicative of regulatory regions for the AS locus on chromosome 1p21, A), and 2q22, B), are shown for heart and aorta tissue samples. Different types of regulatory states are indicated with distinct colours shown at the top of the figure. EnhA (Enhancer Active), EnhW (Enhancer Weak), PromUp/Dn (Chromatin marks characteristic of a promoter region found upstream or downstream of TSS), DNase (DNase, nucleosome-free/open chromatin region), PromP (Promoter poised region, marked simultaneously as active and repressed, poised for activation during development), TssA (Transcription Start Site, Activated), Repr (Repressive marks, heterochromatin). Vertical grey lines indicate the variants found in LD (R^2^ > 0.50) with A) rs7543130 (*) (N=19) or B) rs1830321 (+) (N=50). Variants found to overlap with regulatory regions in any of the five tissues are marked up and indicated as red vertical lines. Long-range chromatin interactions in left ventricle tissue samples are shown for A) the region harboring rs1890753 on chromosome 1p21 with red curved lines, including interactions to promoters for *PALMD, PLPPR4, PLPPR5, DPH5 and SNX7, LOC100129620 and LOC101928270,* and for B) regions harbouring rs13028626, rs6749506, rs2252654, rs4662414, and rs13408842 that directly interact with the promoter regions of *ZEB2, GTDC1, ZEB2-AS1,LINC01412, and TEX41.*

At the *TEX41* locus, five variants in LD with rs1830321 (R^2^ ranging from 0.61 to 1.0) overlapped with regulatory regions (Figure 1-B, upper panel). Chromatin interaction mapping in left ventricular tissue shows that these regulatory regions interact with the promoters of *ZEB2, GTDC1,* and the non-coding RNAs *ZEB2-AS1, TEX41, LINC01412,* and *LOC101928386* (Figure 1-B, lower panel), suggesting that all represent candidate causal genes at the *TEX41* locus.

Of note, *ZEB2* is a DNA-binding transcriptional repressor that interacts with activated SMADs, the transducers of TGFβ signaling. TGFβs are known to play a role in cardiac development and in several aspects of cardiovascular physiology ranging from the cardiomyocyte, to vascular smooth muscle and renal control of blood pressure^29^.

Chromatin interactions between the regulatory regions harboring candidate causal variants at the *PALMD* and *TEX41* loci, were much less frequent in right ventricular tissue and aorta, than the left ventricle, and none overlapped with gene promoters (Supplementary Figure).

In summary, through a large GWAS we discovered two common AS variants on chromosomes 1p21 and 2q22 and replicated the recently reported AS variant in LPA^4^. Assessment of four reported AS variants, including a rare missense variant in *MYH6,* shows associations with BAV, other congenital heart defects, aortic root size and CAD. This implicates cardiac development and atherosclerosis-like processes in the pathophysiology of AS. Finally, our analyses of a genetic risk score based on known CAD variants, suggests that the frequent co-occurrence of AS and CAD is largely explained by genetic predisposition to high Lp(a) or non-HDL-cholesterol.

